# Origin of Cellular Senescence in Ciliates

**DOI:** 10.1101/2024.09.17.613400

**Authors:** Josh Mitteldorf

**Affiliations:** Institute for Frontier Experimental Science

## Abstract

Mixing and sharing of genes is essential for population diversity, which contributes to the resilience and, ultimately, the survival of animal and plant communities. However, sharing of genes is antithetical to individual fitness; hence gene mixing is threatened with extinction if selection rewards selfish (clonal) reproduction. To address this dilemma, mechanisms have evolved to enforce a mandate for gene sharing. In most metazoa, the functions of sex and reproduction are tightly entwined, presumably for the purpose of compelling the sharing of genes. In protists, the functions of sex (conjugation) and reproduction (amitosis) are separate. The mandate for gene sharing is enforced, instead, by a form of cellular senescence. Ciliates can reproduce clonally a few hundred times before they senesce and die. Conjugation resets their biological clock, restarting the cycle of clonal reproduction. The question how reproduction in metazoa came to be linked to sex has been explored in the past, but a fully satisfying account in terms of evolutionary ecology is lacking. The question how senescence in protozoa came to be linked to sex has not been addressed, and is the topic of the current study. I present herein two numerical simulations (IBMs) for the evolution of cellular senescence in ciliates. The results shed light not only on the evolution of senescence in higher life forms, but on more general questions concerning the plausibility of group selection.

## Introduction

Ciliates constitute a phylum of single-celled eukaryotes including *Paramecia, Lacrymaria* and Stentors. In ciliates and some other protists, the functions of replication and sex are separate. Replication is via cell division, cloning with no mixing of genes. Sex occurs via conjugation, which is a forerunner of meiosis. Two individuals exchange cytoplasm and genetic material (including crossover), and replication follows programatically, but not essentially. Two cells enter into conjugation, and two cells emerge; the cells’ identities have become so thoroughly mixed, that it is not meaningful to say which was which originally. Two cells immediately split into four. Conjugation is a slow process compared to clonal reproduction. In paramecia, cloning typically requires minutes, and conjugation requires hours [1].

Ciliate cells have two nuclei, a *micronucleus* that serves as an archival copy of DNA, and a far larger *macronucleus* with polyploidal working copies of each gene. In the micronucleus, genes are arranged on linear chromosomes, but in the macronucleus, there are thousands of separate genes and, typically, hundreds of copies of each distinct gene.

When the cell replicates, the micronucleus undergoes mitosis, while the macronucleus undergoes amitosis [2]. The DNA in the micronucleus separates into two strands, each of which recruits complementary nucleotides to recreate a full and faithful double-stranded copy. But the macronucleus simply splits in two, each daughter macronucleus receiving about half the genes from the parent. This process is called amitosis, to distinguish it from the process by which the micronucleus replicates. Chromosome fragments continue to replicate within the macronucleus to keep pace with the growing cell. But replication in the macronucleus is less faithful than mitosis of the micronucleus. The macronuclear DNA degenerates, and the cell becomes senescent, eventually unable to reproduce.

The cell lineage can be renewed by dissolution of the macronucleus and creation of a new macronucleus from the archival DNA in the micronucleus. But this renewal process is tightly linked to conjugation. Thus, ciliates are required to share genes as a condition for survival of their lineage.

The benefits of gene sharing accrue to the community over long time periods. The benefits of rapid reproduction accrue to the individual immediately. According to classical population genetic theory, there should be no contest – the immediate direct benefit of rapid reproduction should always dominate over the indirect benefit to the community. This was the argument that prevailed in the “group selection” debates of the 1970s [3].

The “masterpiece of nature” [4] is that selfish, unrestrained reproduction has been limited. In the neo-Darwinian model, we think of natural selection rewarding the most efficient replication into the F1 generation. But evolution has also created this barrier, slowing reproduction, for the long-term health of the community. Separation of the sexes and the linkage of sex to reproduction are taken as a starting point in most population genetic analyses and simulations.

In metazoa and protozoa, evolution has tied survival of the lineage to gene sharing. In ciliates, gene sharing is enforced by tying renewal of the macronucleus to conjugation. Without renewing the macronucleus, the macronucleus degenerates and the lineage eventually fails. Without conjugation, a cell cannot renew its macronucleus.

In the models presented here, I attempt to understand the selective dynamic by which cellular senescence evolved in ciliates. How has it come about that renewal of the macronucleus is tied mechanistically to conjugation? This may be regarded as an evolvability adaptation, but a costly one. How has the long-term fitness of the community overcome the short-term fitness cost in this case?

### Relationship to the Diversity Problem

Darwin recognized early that the evolutionary function of natural selection requires a diverse population to be effective, and that the operation of natural selection tends to collapse diversity. He was troubled to the end by the question of how diversity is maintained. The mechanism of Mendelian inheritance, unknown to Darwin at the time, provides a partial explanation; but 150 years on, we still have only partial answers to the question of diversity. Sex and aging are two adaptations that help to maintain population diversity [5-7], but they are both maladaptive for the individual, and the mechanisms by which they evolved are open questions in evolutionary theory.

Why does the community have an interest in assuring that all members share genes? There are two benefits, both related to population diversity. Diversity provides protection against epidemic infections and other threats that can devastate a population that has become too uniformly specialized [8, 9]. And diversity provides fuel for evolutionary change that promises the possibility of better adaptation in the future [10]. Both these are substantial and important benefits, but the first accrues only intermittently and the second only slowly.

### Cellular senescence as an evolutionary adaptation

Senescence first evolved in protists, where the only conceivable benefit was to enforce the imperative to share genes. This contributes to a diverse and viable population, at a substantial cost in individual fitness. To try to understand how the indirect, long-term benefit of cellular senescence can prevail over the direct, short-term cost, I have devised two numerical models, based respectively on the two benefits of population diversity, (1) protection from epidemics, and (2) more rapid pace of population-level adaptation in changing environments. The agent-based models presented here are intended as illustrations only. Our aim is not to promote either of these as the most plausible explanation for how sex came to be obligate in ciliates, but only to offer examples of ways in which the long-term health of the community may overcome a substantial short-term cost to the individual.

### Two Models: Epidemic resistance and Rate of adaptation to change

The first model simulates a subdivided population in which particular genomes are assumed to have a local advantage, so they can reproduce more quickly than all others at a specific location. Countering this, there is the risk of epidemics which preferentially wipe out any genotype that becomes too prevalent locally. After N generations, a lineage is presumed to become sterile, and reaches a dead end, where N is taken in the model to be a gene subject to mutation and selection. When conjugation occurs, the counter is reset for N more replications, and the two genomes cross over to create more diverse offspring. A high value for N means that cell senescence never occurs; an evolved low value represents natural selection of cell senescence.

In the second model, a pool of renewable resource nourishes each subpopulation in a large array of sites, and “fitness” is modeled as efficiency in converting that resource into growth. Reproduction offers an opportunity for mutation, which sometimes increases efficiency. Conjugation offers greater opportunities for increased or decreased efficiency. Conjugation of two individuals of different efficiency expands that differential, creating one individual with much lower efficiency and another with much higher efficiency. The propensity to conjugate is another evolving gene and, another binary gene (the target of our inquiry) determines whether or not the individual keeps count of cell divisions and ceases to replicate after N divisions. (In this second model, N is a parameter for each run, not an evolving gene.) Without cell senescence, the propensity to conjugate evolves ever lower, and it is the sole benefit of cell senescence to prevent this.

By design, these two models are as different as they can be. The first has explicit population division into groups but without explicit geographic structure; the second uses population viscosity on a geographic array. The first makes the assumption that individual cells can sense their own replication count and seek partners for conjugation; in the second, conjugation is a purely random event. In the first, the maximum replication count capacity that is restored upon conjugation is an evolving gene; in the second, it is a fixed parameter of the model. And of course, the two models are based on different fitness benefits of genetic diversity. I don’t intend that either model be taken as a literal account of how cell senescence first evolved, but together the two help to make plausible that natural selection may sometimes favor death over life.

## Results

Both models demonstrate the plausibility of selection for cellular senescence over some regions of parameter space.

### 1) Genetic Diversity and Epidemics

Principal parameters of the model are

- Rate of epidemics
- Rate of conjugation
- Rate of migration among subpopulations

The model is considered to have evolved cell senescence if a steady-state average value (<∞) of the gene for N emerges. (The contrary case is when N continues to evolve higher without limit.)

In Fig 1, we set the rate of migration to zero, and plot the evolved N as a function of the rate of epidemics. The three curves represent three different values for the parameter “rate of conjugation”. Where the curves terminate, no senescence has evolved, i.e., the average N value is evolving higher without limit.

**Fig. 1.**
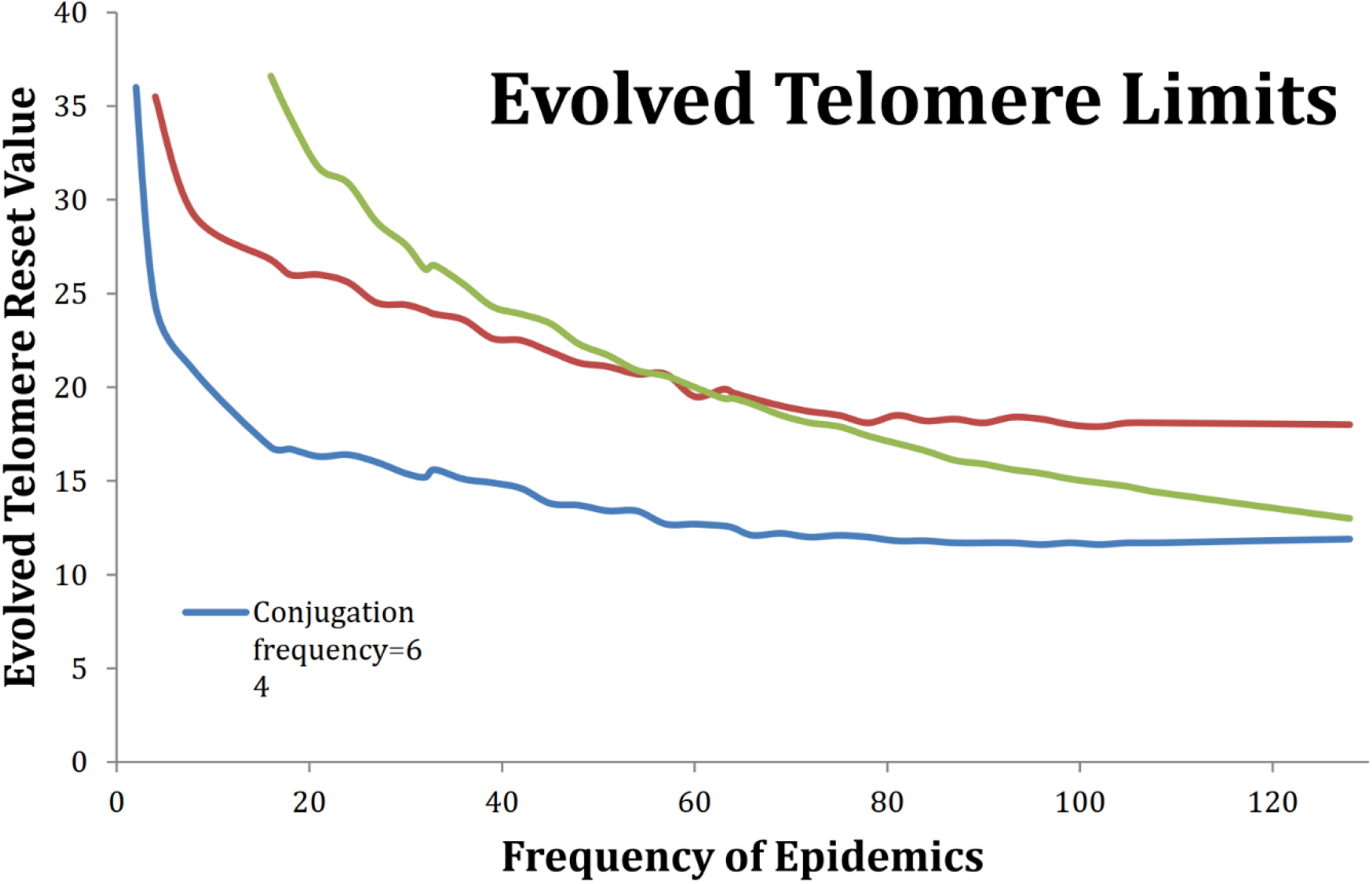

In Fig 2, a portion of 2-dimensional parameter space is mapped. Horizontally across the chart is rate of epidemics, and vertically is rate of conjugation. In each cell, the number indicates the maximum tolerable rate of migration for which senescence is selected. (The maximum was located with a binary search algorithm that ran the simulation repeatedly for different migration parameters, homing in on the maximum.) Zero indicates that cell senescence never evolves.

**Fig. 2.**
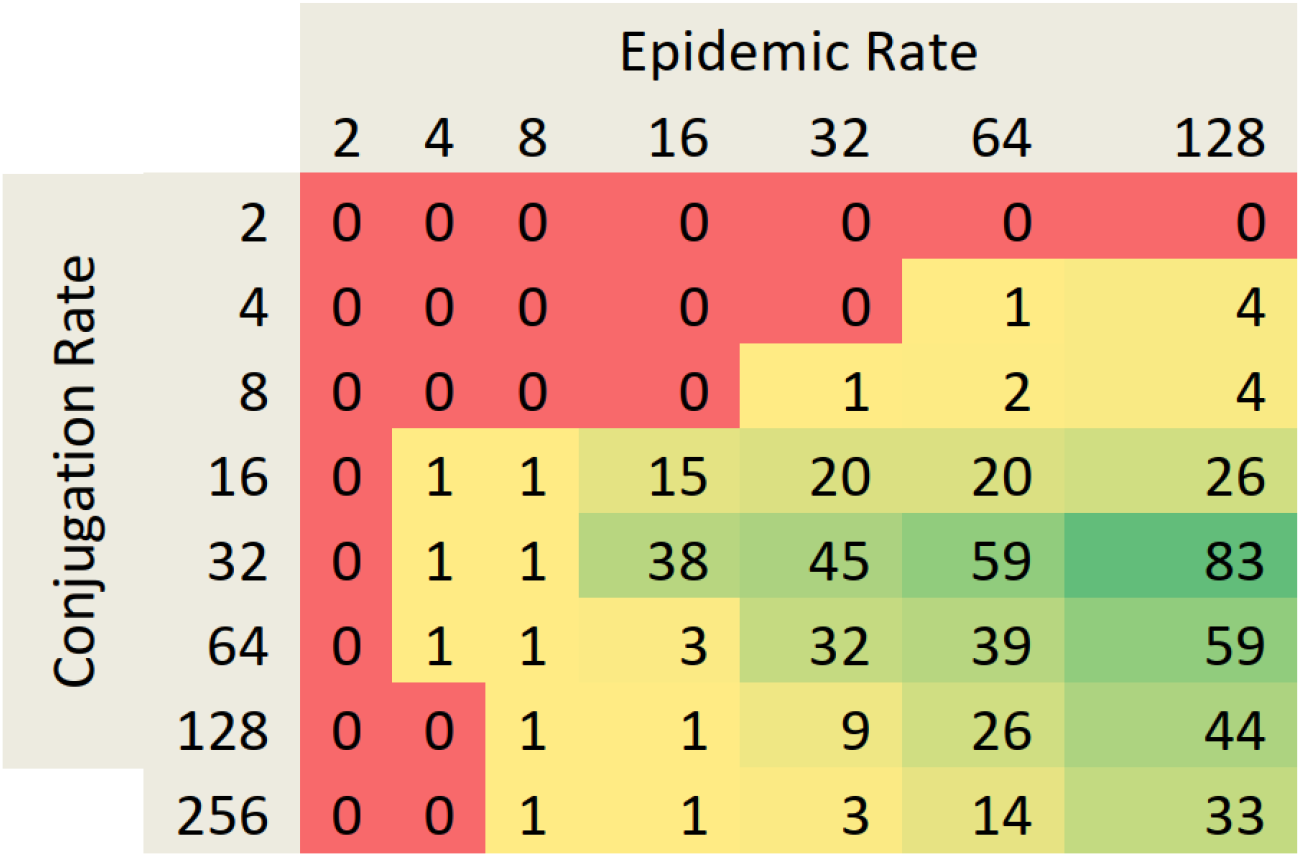

### 2) Rate of evolutionary adaptation

Behavior of this model is influenced most directly by two parameters

- Fitness divergence upon conjugation
- Metabolic cost of conjugation

The results are plotted as a function of these two variables in Fig. 3. Outcomes are somewhat stochastic, and sometimes vary among runs even with identical parameters. The plane defined by the two parameters above is divided with hazy lines into five regions.

A) Non-senescence (unlimited clonal reproduction) evolves to fixation

B) Non-senescence is preferred

C) Coexistence of the senescence and non-senescence

D) Senescence is preferred (limits to clonal reproduction)

E) Senescence evolves to fixation

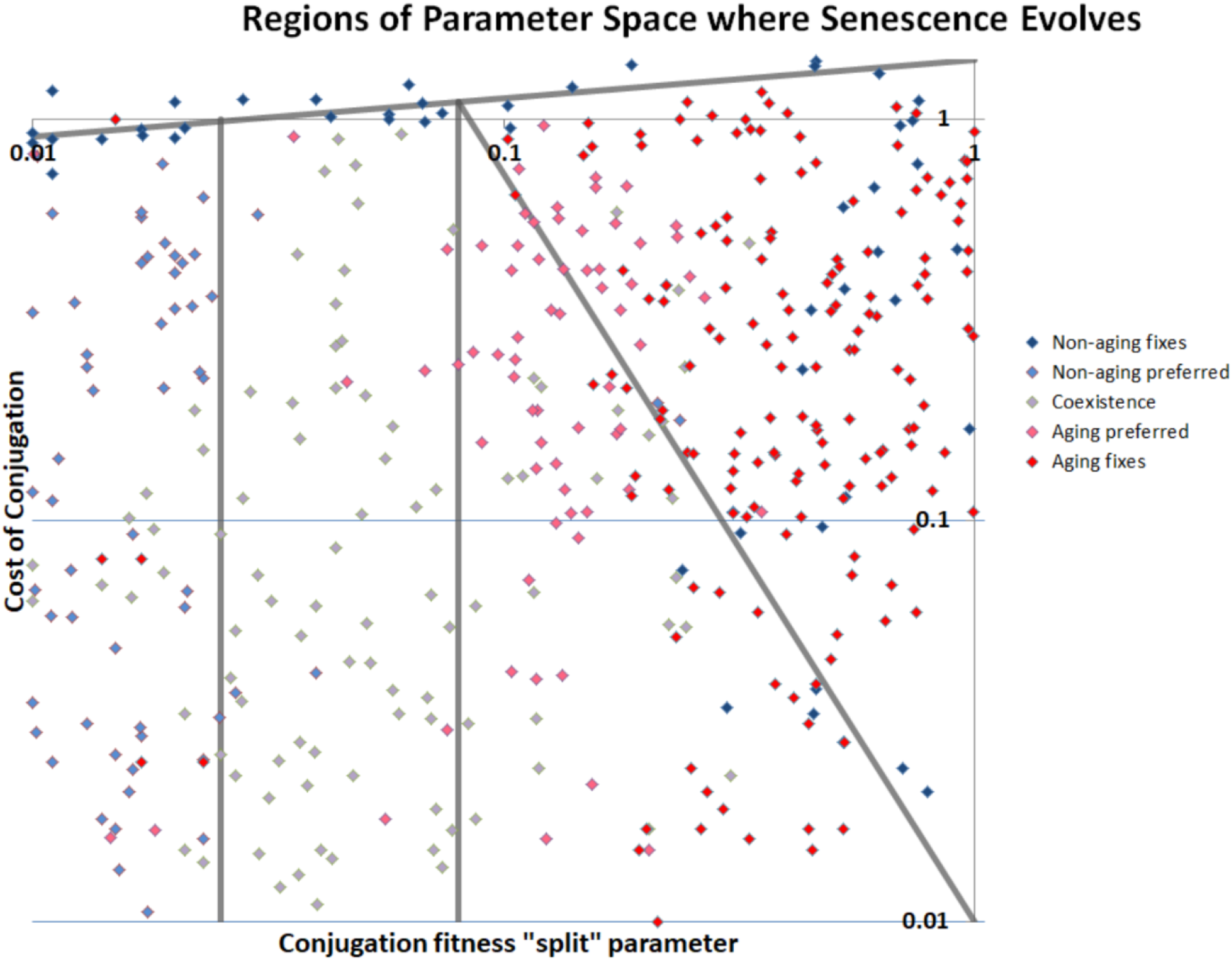

## Discussion

Evolutionary theory is still under the sway of a prejudice that was established in the 1960s and 70s. The prejudice has had ripple effects on gerontological research [11]. Some of our best theorists, including John Maynard Smith, George C. Williams, and William D. Hamilton, established the expectation that group selection has played a negligible role in evolutionary history. They convinced us that all the cooperation we see in the natural world can be understood in an analysis that treated one gene at a time, given only that we must trace the kinship that governed the probability of other cooperating community members bear the same allele.

Dissenting theoretical frameworks of David Sloan Wilson [12] and Michael Gilpin [13] were marginalized [3]. The wise and eloquent naturalist V. C. Wynne-Edwards [14] was denied a place at a table open only to theorists. Eventually, Hamilton came to believe that the theorists’ approach had been too narrow, but he died tragically before his voice could have changed the balance, and his last three volumes were published posthumously [15]. E. O. Wilson [16], though primarily a naturalist, was another adherent of gene-centered evolution until he, too, softened his opposition to group selection later in life [17].

These discussions were excessively theoretical. Some of the strongest examples of altruistic behaviors in nature, including sex, finite lifespans, and ecosystem homeostasis [18] could not be explained by even the expanded mechanistic frameworks of D. S. Wilson [19]

The neo-Darwinian theoretical framework that shaped discussions of group selection was bequeathed to us by R. A. Fisher [20]. Fisher was a mathematician, not a naturalist, and he worked at a time before computer models became feasible. In order to make his model analytically tractable, he made simplifying assumptions. Most relevant, he took sexual reproduction as a given, and he assumed unvarying, steady state populations. It was Gilpin’s insight to allow for exhaustion of prey species, famines, and local extinctions. I have argued that evolution of all forms of altruism are affected by this insight [21].

Evolution has arranged to compel the sharing of genes, and this is remarkable because

- The benefits of gene-sharing are widely-dispersed and accrue only over many generations.
- The individual cost of the adaptation is strong, and it is not borne just by “defectors” that don’t share genes.
- The evolved mechanism of meiosis and its predecessor is metabolically and biochemically complex, necessarily involving a wide range of genetic and epigenetic adaptations.

In this context, the two numerical models presented herein are no more than a beginning toward understanding a phenomenon that challenges our notions of evolution’s mechanics.

### Detailed Model Descriptions

#### 1) Genetic Diversity and Epidemics

A population is subdivided into 128 sites, and there are 128 possible genomes, constructed from 7 genes with two alleles each. At each of the 128 sites, a different one of the 128 genotypes is given a large local advantage, a factor of 2 boost in the probability of reproduction in each time step. (Heuristic: this genotype is highly specialized to the local environment.) The one genotype that fits the local environment quickly grows to dominate the gene pool. But domination of the local site is punished by “epidemics”. With a frequency that is a principal parameter of the model (EPIDEMICFREQ), a random agent is chosen at a random site, and it becomes infected. The infection spreads to all individuals at that one site that share the same genotype as the infected individual, killing them instantly. The greater the prevalence of a particular genotype, the greater the probability that the seed individual who becomes infected will have that genotype. Thus the dominance of any one genotype across a single site poses a risk of local extinction.

Agents have a finite lifespan, with a replication counter limited to N generations of cell division before the cell dies, where N is an evolving gene. The counter is reset to zero with each conjugation. In each time step, there is a constant number of conjugations within each site, where this number (CONJFREQ) is a parameter of the model. Conjugation takes place preferentially among agents with the highest replication counts. (This is an important assumption: cells are presumed to have evolved a means of sensing their senescence, and to have a high propensity for conjugation when the counter is high.) In cases where the gene for N evolves ever higher, senescence is irrelevant. This is interpreted as evolved immortality. On the other hand, if evolution tends to a finite N (fluctuating in steady state), this is interpreted as evolution of cellular senescence.

Conjugation shuffles the 7 genes (bits) from two participating agents and reassigns each gene to either of the two agents at random.

If a site becomes vacant it is always because it has suffered an epidemic after diversity has vanished. The site is repopulated with a seed of six individuals chosen at random from the other 127 sites. Flow of individuals among sites can also take place via random migration, a constant number of individuals exchanged per time step which is a parameter of the model, MIGFREQ. A rate of migration that is sufficiently high makes a subdivided population behave like a single, panmictic population. Therefore, high enough migration always weakens group selection and destroys evolution of cooperation. In asking of the model, “under what circumstances can senescence evolve?”, we have taken the maximum tolerable rate of migration as a measure of the strength of the selective tendency in favor of evolved cell senescence.

#### 2) Rate of Evolutionary Adaptation

The model is realized on a 2D grid, 256*256 grid-cells with toroidal topology. Food is introduced into each grid-cell in each time step, and diffuses into neighboring grid-cells with a diffusion constant that is a parameter of the model. Each grid-cell may support an indeterminate number of individual agents (or none). The model is initialized with one individual per grid-cell, and as evolution progresses, the individuals become more efficient and population increases. In order to keep the model tractable, total population is regulated to be about 10^5^ individuals by downward adjustment (as needed) of the rate that food is supplied. This creates an average of 1.5 individuals per grid-cell, but when efficiency evolves very high at a particular site, the population of that site alone may rise above 10^4^ individuals.

Agents are capable of four behaviors: they may eat, die, reproduce via mitosis, and conjugate with another individual in the same grid-cell.

Individuals have three evolving genes:

##### reproduction-threshold

(a floating point number) How much energy must the individual accumulate before reproducing? This is the primary fitness of the individual. (Lower values correspond to greater fitness.)

##### horniness

(a floating point number) this is the propensity for conjugation. In each time step, every pair of individuals within a cell has a probability of conjugating that is computed as the product of the horniness genes of the two participants.

##### senescing

(two values, 0 or 1) This is the gene for aging. Those individuals with the allele=1 produce daughter cells in each replication with an increased replication counter. Those individuals with allele=0 do not increment their counter, thus they are never subject to senescence.

##### Individuals have two properties that characterize their state

***energy*** is accumulated by eating food, and lost to metabolism and to the energetic cost of conjugation. When energy reaches the *reproduction-threshold*, mitosis takes place.

***remaining cell divisions*** is the second property. Senescing individuals keep track of replications, and die when this number reaches zero; for non-senescing individuals, no replication count is maintained. Senescing individuals emerge from conjugation with a full N cell divisions remaining, where N is another parameter of the model.

In each time step,

- There is a constant probability of death from random mortality.
- Food energy consumed is a constant proportion of the available food concentration in the grid-cell.
- A constant amount of energy is deducted for metabolism as a “cost of living”.
- Mitosis occurs if energy exceeds the *reproduction-threshold*.
- For each pair in a given grid-cell, there is a probability of conjugation, computed as the product of the *horniness* genes of the two participants.

In mitosis, the individual becomes two individuals, each with half the energy. Each of the genes mutates with a small probability. One of the daughters emigrates to a random von Neumann neighbor of the grid-cell of the parent.

In conjugation, two individuals each pay a cost in energy. Genetic exchange is such that the values of the floating-point genes are averaged and then (crucially) one individual emerges with a higher value and the other with a lower value than this average, for each of the two genes. The split between the higher and lower values is another parameter of the model, and this is the payoff for conjugation. One of the two individuals emerges as a loser, with lower primary fitness and is destined for extinction; the other emerges with a primary fitness that is substantially enhanced compared to the range available via mutation alone. The winner thus emerges from conjugation with enhanced potential for evolutionary success. In particular, the primary determinant of fitness is the gene for energy threshold of replication. The winner’s threshold for replication is reduced, so that as available food becomes ever smaller, it is only the rapidly evolving lines that are able to compete.

The most important parameters of the model are:

“split” constant, fitness gained or lost in conjugation

energy cost of conjugation

Other parameters of the model are:

mortality probability per time step

“cost of living” = energy deducted in each time step

diffusion constant for food

N, the max replication count for senescing individuals

proportion of available food that each individual extracts from grid-cell in a time step mutation size during mitosis

(initialization values)

### Why does the model work?

It may seem counterintuitive that the tertiary benefit to fitness from senescence can overcome the direct fitness cost of senescence. One reason that the model succeeds in evolving senescence is that the benefit from conjugation is presumed high. This is embodied in the ratio of the “split” constant to the mutation size. The changes in fitness from mutations are scaled at about 10^−3^ compared to the changes from conjugation. In the model, there are about 30 generations between conjugations, so conjugation has the potential to contribute 1/(30*10^−3^)∼ 30 times as much to the rate of fitness increase as mutation. Another feature of the model dynamic is that the probability of conjugating rises with the square of local population density. When conjugation yields a big boost in fitness locally, there is a surge in reproductive rate, and population in a grid-cell can rise briefly over 1,000, which enhances the opportunities for conjugation 1,000-fold, which leads to a further boost in peak fitness.

## Notes

### Competing Interest Statement

The authors have declared no competing interest.

